# The genetic paradigms of dietary restriction fail to extend life span in *cep-1(gk138)* mutant of *C. elegans* p53 due to possible background mutations

**DOI:** 10.1101/2020.07.09.195503

**Authors:** Anita Goyala, Aiswarya Baruah, Arnab Mukhopadhyay

**Affiliations:** Molecular Aging Laboratory, National Institute of Immunology, Aruna Asaf Ali marg, New Delhi 110067, India; Dept. of Agricultural Biotechnology, Assam Agricultural University, Jorhat 785013, Assam

**Keywords:** *cep-1*, p53, dietary restriction, life span, *cep-1(gk138)*, TJ1

## Abstract

Dietary restriction (DR) increases life span and improves health in most model systems tested, including non-human primates. In *C. elegans*, as in other models, DR leads to reprogramming of metabolism, improvements in mitochondrial health, large changes in gene expression, including increase in expression of cytoprotective genes, better proteostasis etc. Understandably, multiple global transcriptional regulators like transcription factors FOXO/DAF-16, FOXA/PHA-4, HSF1/HSF-1 and NRF2/SKN-1 are important for DR longevity. Considering the wide-ranging effects of p53 on organismal biology, we asked whether the *C. elegans* ortholog, CEP-1 is required for DR-mediated longevity assurance. We employed the widely-used TJ1 strain of *cep-1(gk138)*. We show that *cep-1(gk138)* suppresses the life span extension of two genetic paradigms of DR, but two non-genetic modes of DR remain unaffected in this strain. We find that in *cep-1(gk138)*, two aspects of DR, increased autophagy and the up-regulation of expression of cytoprotective xenobiotic detoxification program (cXDP) genes are dampened. Importantly, we find that background mutation(s) in the strain may be the actual cause for the phenotypic differences that we observed and *cep-1* may not be directly involved in genetic DR-mediated longevity assurance in worms. Identifying these mutation(s) may reveal a novel regulator of longevity required specifically by genetic modes of DR.

## Introduction

Dietary restriction (DR) extends longevity and imparts health benefits to a wide range of metazoans. Research over the past decades, many using *Caenorhabditis elegans*, has revealed the role of evolutionarily conserved transcription factors like FOXA/PHA-4 (Chamoli et al. 2014; Matai et al. 2019; Panowski et al. 2007; Tabrez et al. 2017), NRF2/SKN-1 (Bishop and Guarente 2007), HIF1/HIF-1 (Chen et al. 2009), TFEB/HLH-30 (Lapierre et al. 2013), HNF4/NHR-49 (Chamoli et al. 2014), HSF1/HSF-1 (Steinkraus et al. 2008) and FOXO/DAF-16 (Greer et al. 2007) in the regulation of the transcriptional landscape required to ensure long life span of DR animals. The mechanisms by which DR affects life span are diverse and depends on the mode of implementation. In *C. elegans*, DR may be implemented by genetic means as well as through non-genetic dietary interventions. For example, the *eat-2* mutants that have defective pharyngeal pumping is considered the classical genetic models of DR (Lakowski and Hekimi 1998). We have recently shown that knocking down a serine-threonine kinase gene *drl-1* using RNAi leads to a DR-like phenotype that extends life and health span (Chamoli et al. 2014). Non-genetically, DR can be implemented by diluting bacteria in liquid media (Bishop and Guarente 2007; Panowski et al. 2007) or on solid media plates (Greer et al. 2007; Wu et al. 2019), by peptone restriction (Hosono et al. 1989), by feeding a non-hydrolysable glucose analog (Schulz et al. 2007) or even by complete depletion of bacterial feed (Kaeberlein et al. 2006). Interestingly but quite expectedly, the transcription factor requirements also vary with the model of DR. For e.g., the key transcription factor required in DR regime on solid plates is DAF-16 while for *eat-2* mutants, *drl-1* KD worms and for bacterial dilution in liquid, PHA-4 is required (Chamoli et al. 2014; Greer et al. 2007; Panowski et al. 2007). On the other hand, a DR regime in liquid media on solid support requires SKN-1 (Bishop and Guarente 2007) while increased life span on complete removal of food is dependent on HSF-1 (Kaeberlein et al. 2006). These observations point to the complex modalities of gene regulation brought about by nutrient signalling and DR implemented by various regimes that still need to be elucidated, including identifying new transcriptional regulators.

The p53 protein is a well-known tumour suppressor that has an important role in maintaining genome integrity by inducing DNA repair, cell cycle arrest and apoptosis in response to genotoxic stress (Jolliffe and Derry 2013). Various model organisms like mice (Maier et al. 2004; Tyner et al. 2002), flies (Bauer et al. 2010; Bauer et al. 2007) and worms (Arum and Johnson 2007; Baruah et al. 2014) have been exploited to investigate the role of p53 in the aging process. In *C. elegans*, CEP-1 is the functional ortholog of p53. Interestingly, knocking down *cep-1* by mutation or RNAi leads to a small but significant increase in life span, in a context dependent manner (Arum and Johnson 2007; Baruah et al. 2014). The *cep-1(gk138)* allele has been widely used in studies on *cep-1*. The TJ1 strain was prepared by backcrossing *cep-1(gk138)* ten times with wild-type N2 (Arum and Johnson 2007) and is available from the Caenorhabditis Genetics Center (CGC). In this study, we asked whether TJ1 *cep-1(gk138)* produced life span extension when grown under different DR regimes. We found that the genetic modes of DR, namely *drl-1* KD and *eat-2* mutant, failed to extend life span in TJ1. However, non-genetic DR regimes, namely BDR and 2-DOG were unaffected. We show that under genetic DR, two important cellular processes, autophagy induction and cytoprotective response, through the activation of the xenobiotic detoxification program (cXDP) gene expression, fail to get upregulated in TJ1 *cep-1(gk138)*. Also, TJ1 differentially affects the two genetic models of DR in terms of regulation of fat storage. However, we found that after backcrossing the TJ1 allele 2 times, or on using two other mutant alleles, life span extension on *drl-1* RNAi became independent of the absence of *cep-1*, suggesting that background mutation(s) in the TJ1 strain may suppress genetic DR-mediated longevity extension. Interestingly, *cep-1(gk138)* TJ1 as well as the 12X backcrossed mutant consistently enhanced dauer formation in the insulin signalling defective *daf-2(e1370)*, suggesting that it may be a *bona fide* property of *cep-1.* Since TJ1 differentially influences the two genetic models of DR, it will be interesting to identify the background mutation(s).

## Results

### *Cep-1(gk138)* (TJ1) suppresses life span of genetic paradigms of DR

The *gk138* allele (TJ1; 10x backcrossed) was obtained from the Caenorhabditis Genetics Center (CGC) and has a 1660 bp deletion in the *cep-1* gene. In our lab, we are characterizing a genetic model of DR where knocking *drl-1* kinase gene leads to a DR-like phenotype, resulting in a dramatic increase in life span (Chamoli et al. 2014). We found that *drl-1* KD worms failed to show life span extension in TJ1 *cep-1(gk138)* (**Figure 1A**) [to be called *cep-1(gk138)*]. The *eat-2* mutants represent the classical genetic models of DR (Lakowski and Hekimi 1998). We compared the life span of *eat-2(ad1116)* with the long-lived *eat-2(ad1116)*;*cep-1(gk138)* and found that the life span of the former is completely suppressed (**Figure 1B**). Similar results were observed with another allele, the *eat-2(ad465)* (**Figure 1C**). Together, we found that TJ1 *cep-1(gk138)* mutant suppresses life spans of two genetic paradigms of DR, the *eat-2* mutants as well as the worms grown on *drl-1* RNAi.

**Figure 1.**
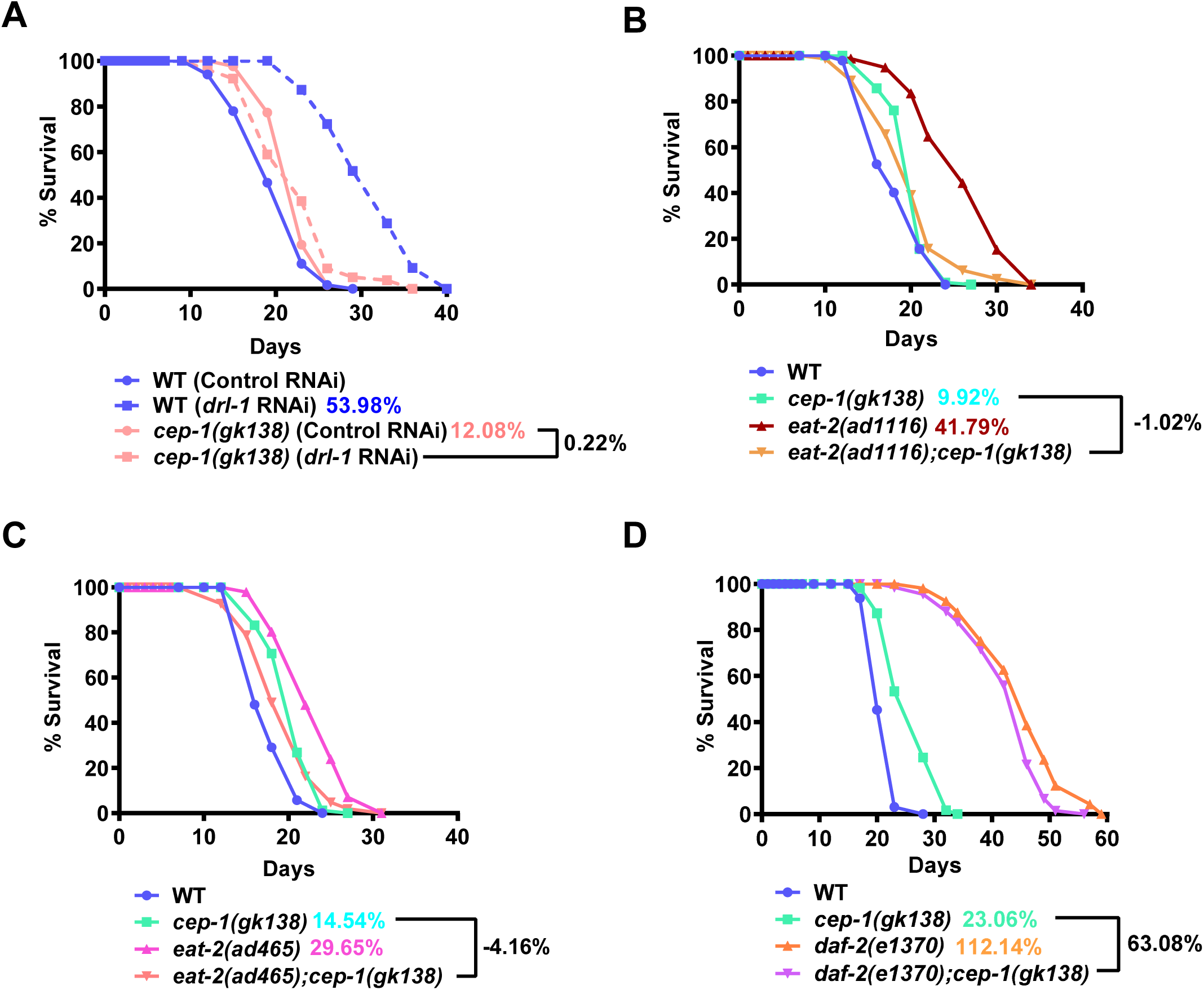
**(Also see Figure S1). In *cep-1(gk138)*, genetic paradigms of Dietary Restriction fail to extend life span. (A)** The life span extension upon *drl-1* KD in WT is suppressed when *cep-1(gk138)* is used. **(B)** The extended life span of *eat-2(ad1116)* is supressed in *eat-2(ad1116);cep-1(gk138).* **(C)** The extended life span of *eat-2(ad465)* is supressed in *eat-2(ad465);cep-1(gk138).* **(D)** The life span of *daf-2(e1370)* is partially suppressed when combined with *cep-1(gk138)*, as in *daf-2(e1370);cep-1(gk138)*. Life spans were performed at 20 °C. Details of life span are provided in Table S1.

We next asked whether other longevity pathways are influenced by *cep-1(gk138)*. We analysed the effect on the long life span of the reduced Insulin/IGF-1 signalling (IIS) mutant, *daf-2(e1370)*. The *daf-2(e1370);cep-1(gk138)* showed a partial reduction compared to the life span of *daf-2(e1370)* (**Figure 1D**), showing that the effect was more robust in case of DR. Interestingly however, the dauer formation of *daf-2(e1370)* was dramatically enhanced in *daf-2(e1370);cep-1(gk138)* (**Figure S1**). This shows that life span and dauer development is differentially regulated in the IIS pathway mutant in combination with TJ1 *cep-1(gk138).*

DR may be initiated non-genetically in worms either by diluting the bacterial feed (BDR) (Panowski et al. 2007) or by using a non-hydrolysable glucose analog, 2-deoxyglucose (2-DOG) (Schulz et al. 2007). We asked if the *cep-1(gk138)* disrupts the life span extension of the non-genetic paradigms of DR as well. We observed that bacterial dilution also generated the typical bell-shaped curve in the BDR assay of *cep-1(gk138)*, as seen in WT worms, when average life spans were plotted against the bacterial dilutions (**Figure 2A**). Likewise, on supplementation of 2-DOG, *cep-1(gk138)* showed life span extension similar to WT worms (**Figure 2B**). Overall, both the non-genetic DR paradigms inTJ1 *cep-1(gk138)* could increase life span indicating that it only affects the genetic modes of DR.

**Figure 2.**
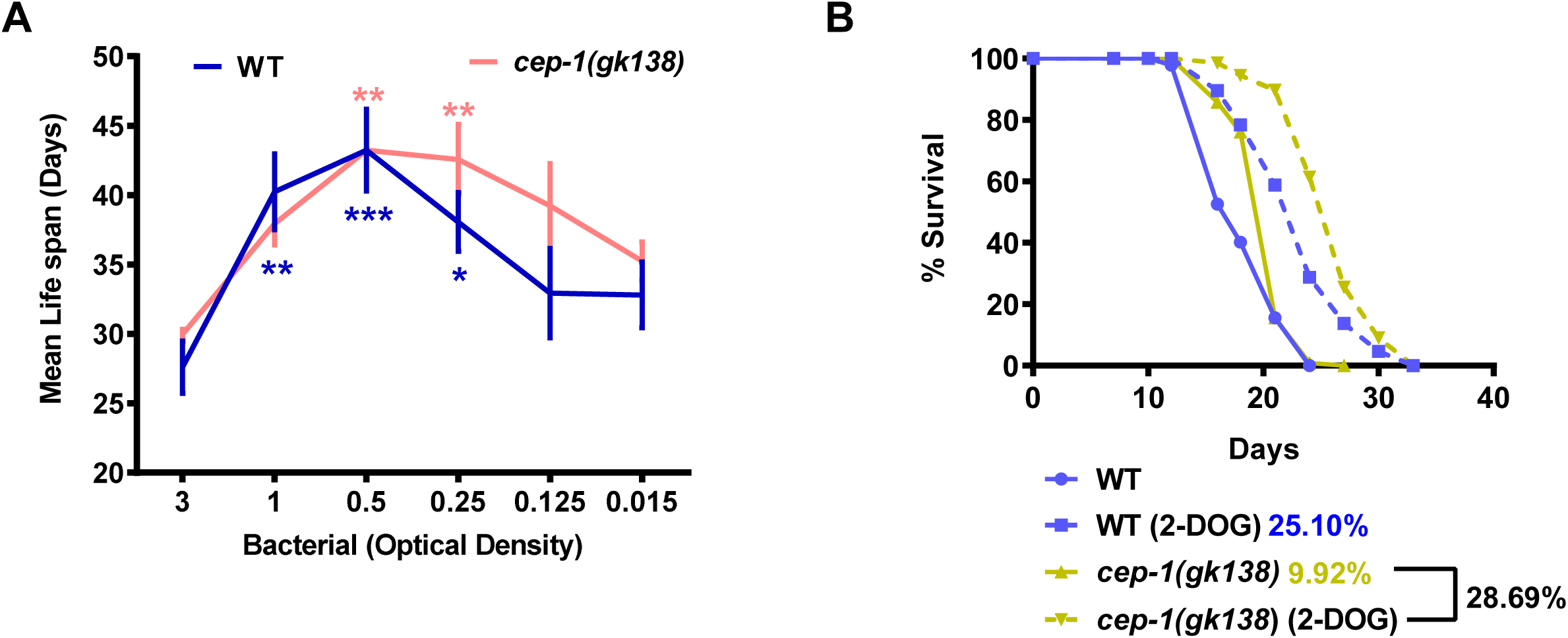
**The life span extension on non-genetic modes of DR remain unaffected in *cep-1(gk138).* (A)** A bell-shaped curve is observed when WT or *cep-1(gk138)* worms are grown on different dilutions of bacterial feed and the average life span plotted against the OD_600_. Average of three biological replicates is shown. Error bars represent SEM. Statistical analysis was performed using two-way Annova. \**P*≤0.05, \*\**P*≤0.01, **\*\***\**P*≤0.001. **(B)** Supplementation of 2-DOG extends life span in *cep-1(gk138)* mutant similar to WT. Life span was performed at 20^°^C and details are provided in Table S1.

### Autophagy fails to be upregulated in *cep-1(gk318)* (TJ1) undergoing DR

A prominent marker for long-lived mutants is increased autophagy. DR-induced autophagy requires the transcription factors FOXA/PHA-4 and HLH-30, implying that autophagy is transcriptionally regulated during DR (Hansen et al. 2008; Lapierre et al. 2013). We asked whether autophagy is mis-regulated in the TJ1 *cep-1(gk138)* undergoing the genetic paradigms of DR, providing a possible reason for life span suppression. We observed that the increase in autophagosome number, as determined by the number of puncta in the seam cells of the L3 stage *lgg-1::gfp* transgenic worms, after knocking down *drl-1* in WT background, was completely suppressed in *cep-1(gk138)* (**Figure 3A, S2A**). Next, we found that the increased puncta in *eat-2(ad1116);lgg-1::gfp* was also suppressed in *eat-2(ad1116);cep-1(gk138);lgg-1::gfp* (**Figure 3B, S2B**). Basal levels of autophagy were unchanged in *cep-1(gk138)*, compared to WT. This implies that *cep-1(gk138)* influences autophagy in the two genetic models of DR.

**Figure 3.**
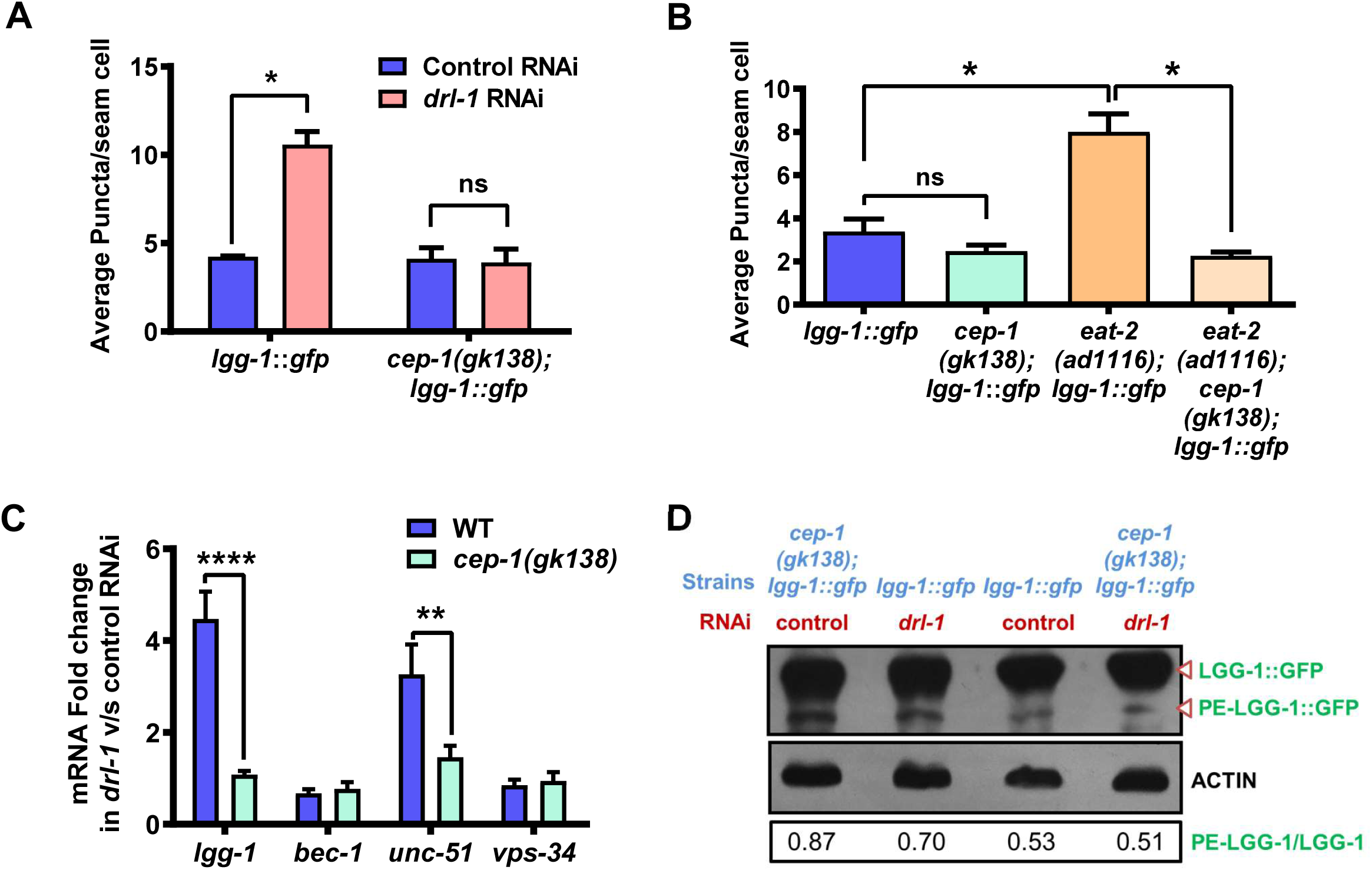
**(Also see Figure S2). *Cep-1(gk138)* regulates autophagy during DR. (A)** Increased autophagosome formation in *lgg-1::gfp* grown on *drl-1* RNAi is suppressed in *cep-1(gk138); lgg-1::gfp*. **(B)** The increased autophagosome formation in *eat-2(ad1116)* is suppressed in *eat-2(ad1116);cep-1(gk138).* Quantification of GFP puncta averaged over 2 biological repeats is shown. Error bars represent SEM. No. of animals analysed, n≥15. Unpaired two-tailed t-test. \**P*≤0.05, ns= non-significant. **(C)** Transcript levels of autophagy genes, *lgg-1, bec-1, unc-51, vps-34* in WT and *cep-1(gk138)* on control and *drl-1* RNAi. Average of at least three biological replicates is shown. Error bars represent SEM. Two-way Annova was used for statistical analysis. \*\**P*≤0.01, \*\*\*\**P*≤0.0001. **(D)** Western blot using anti-GFP antibody to detect the unmodified LGG-1 or PE-LGG-1. The *lgg-1::gfp* or *cep-1(gk138);lgg-1::gfp* worms were grown on control or *drl-1* RNAi. Densitometric quantification of protein bands was done using ImageJ and ratio of PE-LGG-1/LGG-1 is shown. β-ACTIN was used as a loading control. One representative blot out of the two biological replicates is shown.

We next followed up this observation by determining the expression levels of four of the important autophagy genes (*lgg-1, bec-1, vps-34*, and *unc-54*) under control and *drl-1* KD conditions, using wild-type and *cep-1(gk138)*. We observed a transcriptional up-regulation of *lgg-1* and *vps-34* on knock-down of *drl-1*as compared to control RNAi, and this increase was reduced in *cep-1(gk138)* (**Figure 3C**). To biochemically determine the level of autophagy up-regulation, the extent of PE-LGG-1 formation was evaluated using western blot analysis. We observed that on *drl-1* KD, there was an increase in the PE-LGG-1 band intensity, corresponding to the increased autophagosome numbers seen in the puncta assay. In line with the autophagosome assay, the PE-LGG-1 band intensity reduced in *cep-1(gk138)* as compared to wild-type on *drl-1* KD (**Figure 3D**).

In order to find whether the regulation of autophagy in *cep-1(gk138)* was specific to the modes of DR used in this study, we determined whether autophagy induction in *daf-2(e1370)* is also influenced by the TJ1 *cep-1(gk138)*. In agreement with previous data (Melendez et al. 2003), we also found that reduced IIS led to increased autophagic puncta. However, this increase was maintained in the *cep-1(gk138)* background (**Figure S2C, D, E**). These observations show that TJ1 *cep-1(gk138)* regulates autophagy in response to genetic modes of DR, but not on lowering IIS signalling.

### Fat storage is differentially affected in the two genetic DR models by *cep-1(gk138)*

One of the hallmarks of DR is metabolic reprogramming towards increased fatty acid oxidation that leads to depletion of stored fat (Chamoli et al. 2014). We asked whether *cep-1(gk138)* would have differences in depletion of stored fat in the two genetic models of DR. We found that on *drl-1* KD in both WT and TJ1 *cep-1(gk138)*, fats stores were depleted, as measured by Oil Red O staining of fixed worms (**Figure 4A**). Surprisingly, in the *eat-2(ad1116);cep-1(gk138)*, the reduced fat stores of *eat-2(ad1116)* was partially restored (**Figure 4B**). Similar observations were made for *eat-2(ad465);cep-1(gk138)* (**Figure S3**). Together, TJ1 *cep-1(gk138)* responds differently to the two genetic modes of DR.

**Figure 4.**
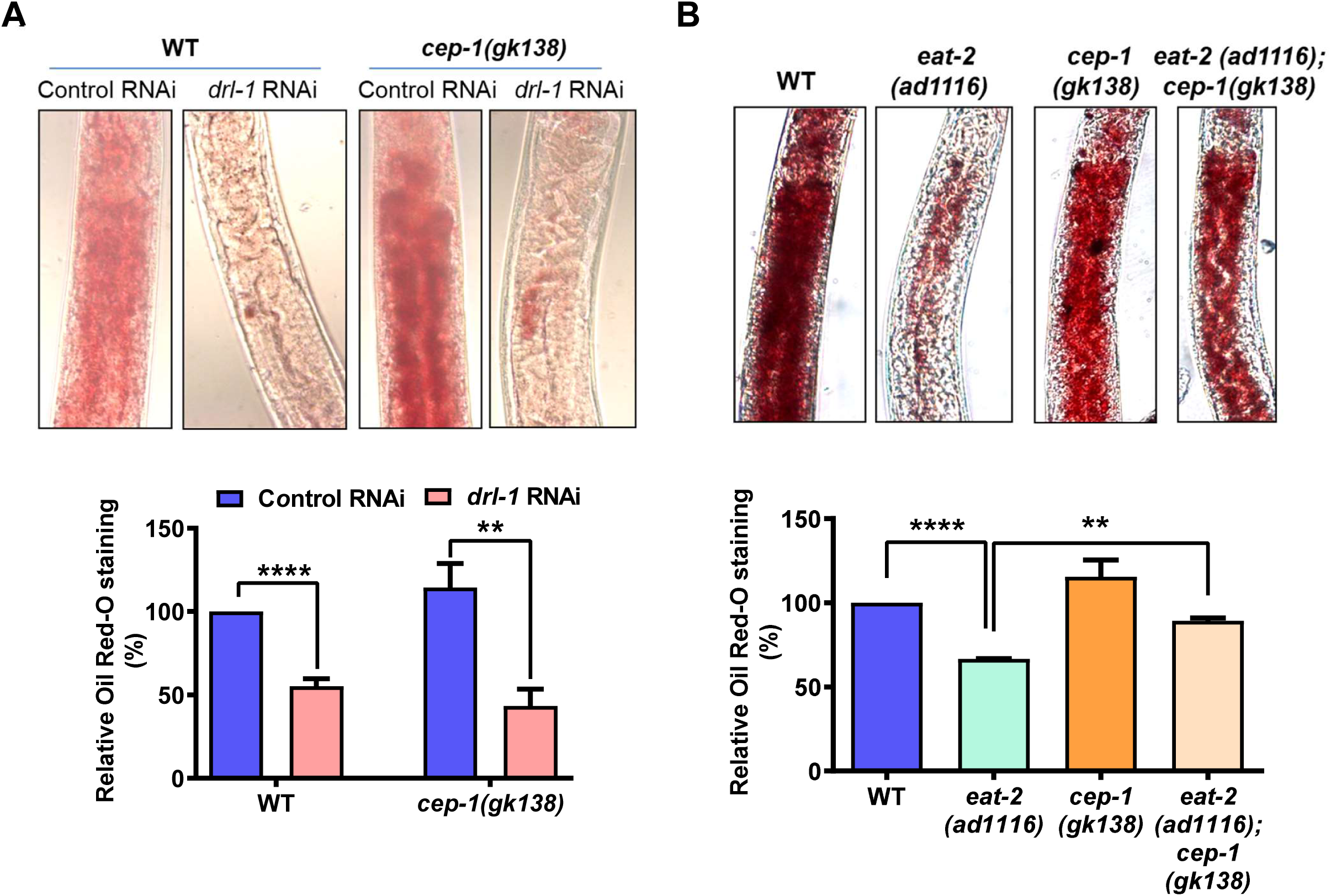
**(Also see Figure S3). Fat storage is differentially regulated by *cep-1(gk138)*. (A)** Oil Red O staining of WT or *cep-1(gk138)* grown on control or *drl-1* RNAi. Lowering of fat storage takes place in both the strains grown on *drl-1* RNAi. **(B)** Oil Red O staining of *eat-2(ad1116)* or *eat-2(ad1116);cep-1(gk138)*. Lowering of fat storage in *eat-2(ad1116)* is attenuated in *eat-2(ad1116);cep-1(gk138)*. Images were captured at 100X magnification. Quantification averaged over at least two biological replicates is shown below in each case. Error bars represent SEM. Unpaired two-tailed *t*-test. \*\**P*≤0.01, \*\*\*\**P*≤0.0001.

### Suppression of cytoprotective gene activation in *cep-1(gk138)* undergoing genetic DR

Previous study from our lab has shown that *drl-1* KD leads to up regulation of the cytoprotective xenobiotic detoxification pathway (cXDP) genes to ensure DR-mediated life span enhancement (Chamoli et al. 2014). This up-regulation requires the conserved transcription factors like PHA-4, NHR-8 and AHR-1. Since CEP-1 is also known to regulate detoxification genes like *gst-4* (Leung et al. 2012), we asked whether the cXDP genes are optimally upregulated in TJ1 *cep-1(gk138)* undergoing genetic modes of DR. Quantitative RT-PCR showed that the phase I and phase II cXDP were up-regulated in wild-type on *drl-1* KD. However, the extent of up regulation of some of these genes, like *cyp-33, cyp-35, cyp-37* and *ugt-16* was significantly reduced in *cep-1(gk138)* grown on *drl-1* RNAi. Other genes, like *cyp-32* and *gst-28* were downregulated but not significantly (**Figure 5A**). The transcript levels of three of these genes were also determined in *eat-2* mutant and two were found to be reduced in the TJ1 background (**Figure 5B**).

**Figure 5.**
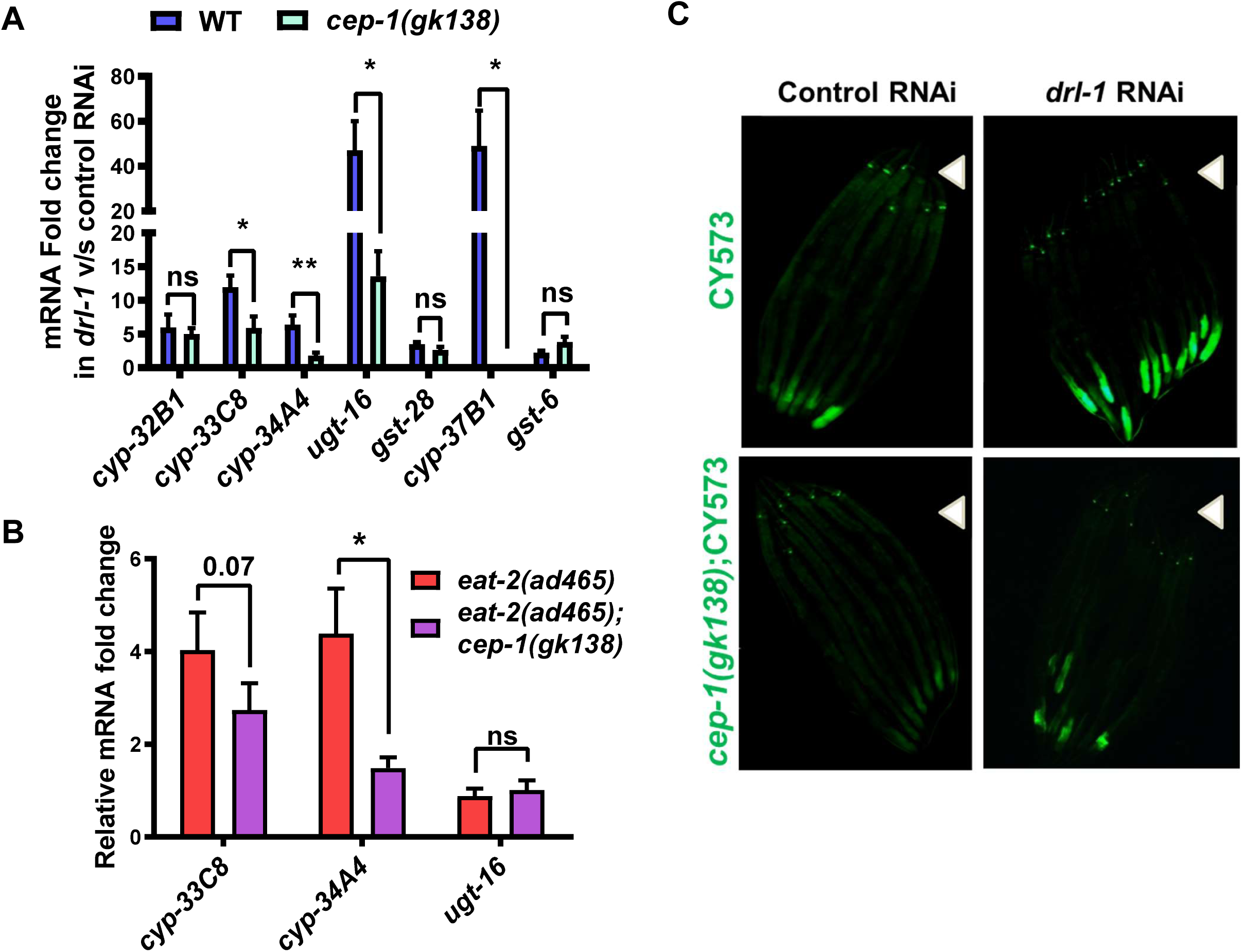
**The increased expression of cXDP genes in *cep-1(gk138)*, undergoing genetic DR, is partially attenuated. (A)** The mRNA levels of cXDP genes are increased in WT grown on *drl-1* RNAi, but partially suppressed in *cep-1(gk138)*. Average of at least three biological replicates is shown. Error bars represent SEM. Unpaired two-tailed *t*-test. \**P*≤0.05, \*\**P*≤0.01, ns=not significant. **(B)** The increased levels of cXDP genes in *eat-2(ad465)* is reduced in *eat-2(ad465);cep-1(gk138)*. Average of at least three biological replicates is shown. Error bars represent SEM. Unpaired two-tailed *t*-test. \**P*≤0.05, ns=not significant. **(C)** Representative images showing increase in *pcyp-35B1:gfp* expression on *drl-1* KD in WT, which abrogated in *cep-1(gk138)*. n≥20. Images were captured at 100X. Arrow heads point towards the head of the worms.

We also used the transcriptional reporter-containing transgenic strain, *Pcyp-35B1::gfp* that has the promoter of *Cyp-35B1*, a gene of phase I XDP, driving the expression of GFP in the intestine. When *drl-1* is knocked down in *Pcyp-35b1::gfp*, the expression of GFP increases but this was suppressed in *cep-1(gk138);Pcyp35b1::gfp* (**Figure 5C**). Together, these results show that the optimal expression of cXDP genes during DR is hampered in TJ1 *cep-1(gk138)*.

### Background mutation(s) in TJ1 *cep-1(gk138)* may be responsible for suppressing DR life span

The TJ1 strain obtained from CGC is 10x backcrossed. We observed that while the original 10x (**Figure 1A**) as well as the 11x backcrossed [the CGC TJ1 *cep-1(gk138)* backcrossed once] strains showed complete suppression of life span when grown on *drl-1* RNAi (**Figure 6A**), the 12X strain [the CGC TJ1 *cep-1(gk138)* backcrossed twice] failed to do so (**Figure 6B**). We reordered the stains from CGC along with VC172 that is 0X backcrossed. We found that the newly acquired TJ1 strain still led to complete suppression of *drl-1* RNAi life span extension (**Figure 6C**) while similar observation was not seen with the VC172 (**Figure 6D**). Further, we tested two other alleles of *cep-1*, namely *cep-1(ep347)* and *cep-1(lg12501)* as well as another 12X backcrossed line generated earlier (Baruah et al. 2014) and found that life span was not suppressed as before (**Figure 6E, F, S4A**). Together, we believe that a background mutation(s) in the 10X TJ1 strain of *cep-1(gk138)* suppresses the positive effects of the two different genetic modes of DR. Interestingly though, the dauer enhancement phenotype of *daf-2(e1370)* is still retained in the *daf-2(e1370);cep-1(gk138)*(12X) **(Figure S4B)**, suggesting either that this phenotype is specific to *cep-1* or that the mutation is closely linked to *daf-2* locus. Future studies to characterize and identifying the mutation(s) may lead to finding novel regulators of DR life span.

**Figure 6.**
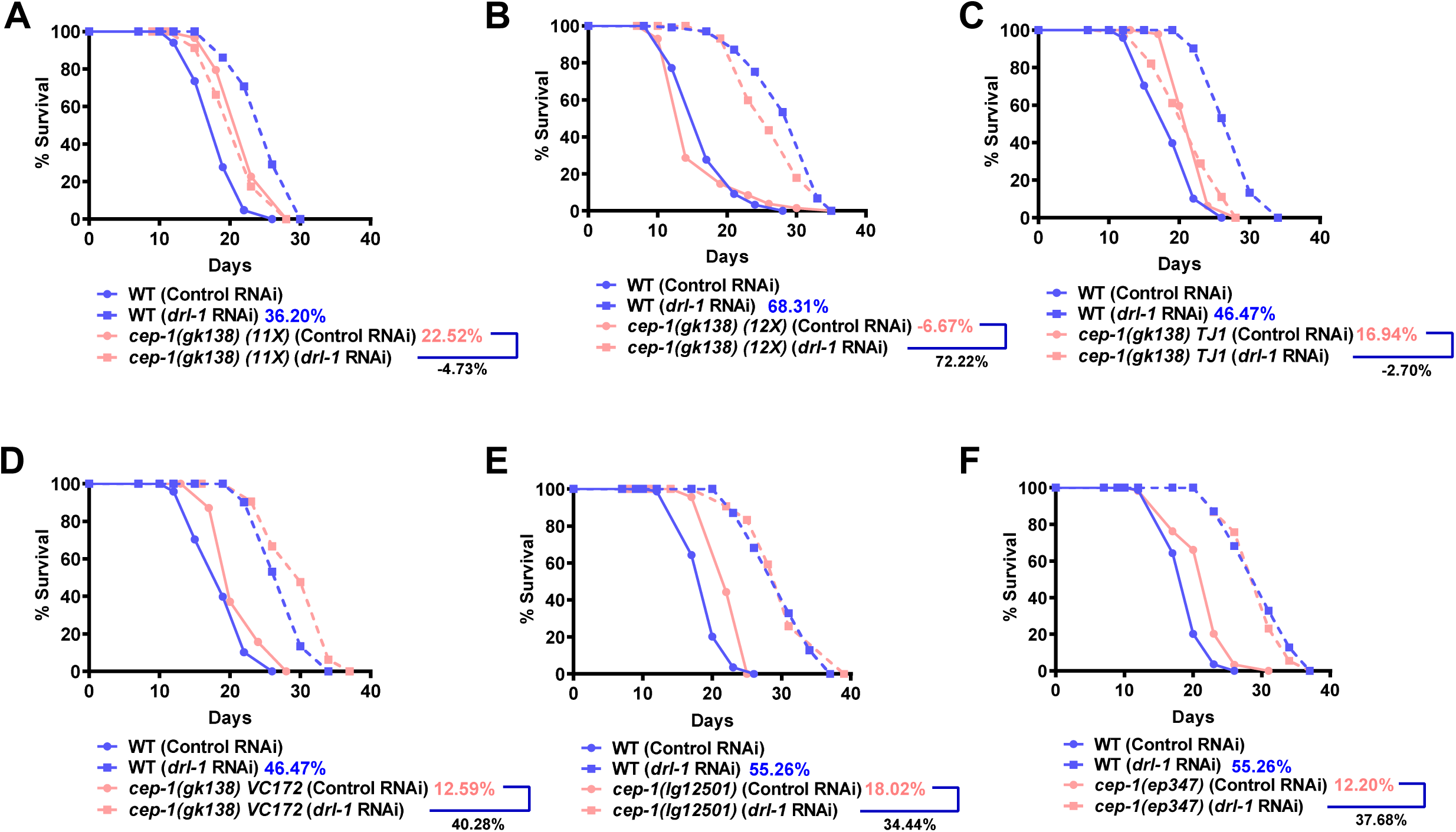
**(Also see Figure S4). Background mutation(s) in *cep-1(gk138)* may be responsible for suppressing life span of genetic paradigms of DR. (A)** Life span analysis of TJ1 *cep-1(gk138)* 11X backcrossed strain, **(B)** 12X backcrossed strain, **(C)** TJ1 strain reordered from CGC, **(D)** VC172 *cep-1(gk138)* 0X backcrossed strain, **(E)** *cep-1(lg12501)* and **(F)** *cep-1(ep347)* along with WT, grown on control or *drl-1* RNAi. Life span was performed at 20^°^C and details are provided in Table S1.

## Discussion

P53 is a multi-functional transcription factor that regulates various biological processes such as DNA repair, apoptosis, cellular senescence, autophagy etc. in response to diverse stresses like DNA damage, oxidative stress, and nutrient deprivation. In this study, we investigated the role of the *C. elegans* ortholog of p53, *cep-1*, in DR-mediated longevity assurance. We employed the commonly-used allele, TJ1 *cep-1(gk138)* that is 10X backcrossed and available from CGC. Although we found that the strain prevented genetic modes of DR from increasing life span, the effect may be the result of background mutation(s).

The TJ1 strain has been widely used to study the role of *cep-1* in aging in *C. elegans*. One of the first such studies showed that the TJ1 strain has a significantly enhanced life span that is dependent on the FOXO transcription factor DAF-16 (Arum and Johnson 2007). Later, the same allele was used to show that the increased life span was dependent on functional autophagy (Tavernarakis et al. 2008). Ventura et al., also used TJ1 to show that *cep-1* mediates the life span effects of the mitochondrial mutants, depending on the mitochondrial bioenergetic stress (Ventura et al. 2009). In the TJ1 strain, mild genetic mitochondrial perturbations that are known to increase life span failed to do so. Additionally, the life span shortening effects of severe mitochondrial disruption also required *cep-1* (Ventura et al. 2009). The TJ1 strain was also used in a later study which further characterized the opposing effects of *cep-1* on life span of mitochondrial mutants (Baruah et al. 2014). However, in that study the TJ1 strain was backcrossed two times with wild-type.

P53 is a complex biological molecule that has both pro-as well as anti-aging properties that depend on the physiological context (de Keizer et al. 2010). It is the dominant tumour suppressor that facilitates DNA repair by halting cell cycle. It also directly impacts the activity of DNA repair systems (Williams and Schumacher 2016). Understandably, loss of p53 would lead to accumulation of mutations in the DNA. The TJ1 *cep-1(gk138)* allele may have accumulated unlinked mutations due to the absence of the p53 ortholog. As a result, the strain may show additional phenotypes that are not linked to *cep-1*. Although *Drosophila* studies indicate that DmP53 may increase life span in a manner similar to caloric restriction (Bauer et al. 2010; Bauer et al. 2005), our study in *C. elegans* suggests that it may be due to background mutation(s). Considering our results, the role of p53 in DR may not have evolved in *C. elegans* or other nematodes. In future, one has to carefully design experiments to decouple the role of background mutation(s) in the highly mutable strain lacking p53, to decipher the real biological function of this important transcription factor on longevity. It will be prudent to experiment with additionally backcrossed TJ1 as well as validating the outcomes parallelly in other alleles like *cep-1(lg12501)* and *cep-1(ep347)*.

### Experimental procedures

#### Strains

All the strains were maintained at 20°C, unless otherwise stated, on a lawn of *E. coli* (OP50) bacteria seeded on standard Nematode Growth Media (NGM) plates (Stiernagle 2006).

**Strains used in the study are**: N2 Bristol as wild-type, TJ1 *cep-1(gk138) I, eat-2(ad1116) II, eat-2(ad465) II, daf-2(e1370) II, adIs2122 [lgg-1p::GFP::lgg-1 + rol-6(su1006)]*, CY573-*bvIs5 [cyp-35B1p::GFP + gcy-7p::GFP]*, XY1054 *cep-1(lg12501) I*, VC172 *cep-1(gk138)* I, CE1255 *cep-1(ep347).*

**Double mutants generated in the study are**: *eat-2(ad1116);cep-1(gk138),eat-2(ad465); cep-1(gk138), daf-2(e1370);cep-1(gk138), cep-1(gk138);adIs2122, eat-(ad1116); cep-1(gk138);adIs2122, eat-(ad1116);adIs2122, daf-2(e1370); cep-1(gk138);adIs2122, daf-2(e1370);adIs2122, cep-1(gk138);bvIs5* [named *cep-1(gk138);CY573*].

#### Preparation of RNAi plates

Nematode Growth Media (NGM) was supplemented with 100 μg/ml ampicillin and 2 mM IPTG to prepare RNAi plates. *E. coli* HT115 was transformed with the L4440 plasmid or *drl-1* cloned in L4440 plasmid. A single colony of bacteria was grown in Luria Bertani (LB) broth containing 100 μg/ml Ampicillin and 12.5 μg/ml Tetracycline, overnight at 37 °C in a shaker incubator. Next day, this culture was used as primary inoculum for sub-culturing in a fresh batch of LB media containing 100 μg/ml ampicillin at a ratio of 1:100 and grown until OD_600_ reached 0.6 at 37 °C. The bacterial cells were then harvested at 5000 rpm, 4 °C and resuspended in 1X M9 buffer (at a dilution of 1:10) containing 100 μg/ml Ampicillin and 1 mM IPTG. This bacterial suspension was then seeded onto the RNAi plates and dried at room temperature for 2 days before use.

#### Life span Assay

Gravid adult hermaphrodite worms were bleached, washed and the eggs collected by centrifugation. These eggs were allowed to hatch on plates containing the OP50 feed or RNAi bacteria. When they reached young adult stage, they were transferred to NGM OP50 or RNAi plates, overlaid with FUDR (final concentration of 0.1 mg per ml). Worms were scored as dead or alive by prodding them with a platinum wire every 2-3 days. Unhealthy worms or worms that crawled to the sides of the plates were censored from the population. Life span graph was plotted as percentage survival on Y axis and the number of days on the X axis. Life spans are expressed as average life span ± SEM for all the life span experiments. Life span data summary is reported in **Table S1**.

#### BDR Life span Assay

BDR was performed as published earlier (Chamoli et al. 2014; Panowski et al. 2007). Briefly, *E. coli* HT115 with L4440 plasmid (referred to as control RNAi) was streaked on a LB agar plate, supplemented with Ampicillin (100 μg per ml) and Tetracycline (12.5 μg per ml) and incubated at 37 °C for 14-16 hours. A single colony was chosen from this plate and inoculated into 200 ml LB containing Ampicillin (100 μg per ml) in a 2 litre flask and grown at 37 °C for 12 hours in an incubator shaker. The bacterial cells were pelleted by centrifugation at 5000 rpm, 4 °C for 10 minutes, which was then resuspended in S-basal-cholesterol-antibiotics solution (cholesterol 5 μg per ml, Carbenicillin 50 μg per ml, Tetracycline 1 μg per ml, Kanamycin 10 μg per ml) supplemented with 2 mM IPTG. This was then diluted to the required optical density (OD_600_) using S-basal-cholesterol antibiotics solution, forming different dilutions of bacteria which were kept at 4°C for a maximum of 2 weeks.

Gravid adult worms maintained on OP50 bacterial feed were bleached and eggs were kept on a 60 mm control RNAi-seeded NGM RNAi plate. On reaching young adult stage, FUDR (100 μg per ml) was added to each plate to prevent progeny development. After about 24 hours, approximately 10-15 worms were transferred to each well of a 12-well plate containing 1 ml of S-basal-cholesterol-antibiotics solution with FUDR (100 μg per ml). The plate was kept on a shaker for 1h to remove adhering bacteria. Meanwhile, the diluted bacterial suspensions were aliquot to separate 12-well cell culture plates, 1 ml solution per well along with FUDR at 100 μg per ml. After the worms were washed off the bacteria, 10-12 worms were moved from the S-Basal to the diluted bacterial suspension with the help of a glass pipette connected to a P200 pipette. Every 3-4 days, worms were moved to fresh bacterial solutions. Before this transfer, they were scored for movement by prodding using a platinum wire. FUDR supplementation in diluted bacterial suspension was only required for the first 8 days. Worms that did not respond to gentle prodding with a worm pick were scored as dead and removed. During experiments, the temperature was maintained at 20°C with continuous rotation of plates at 100rpm in an incubator shaker (Innova 42 incubator shaker, New Brunswick Scientific, New Jersey, USA). Bacterial dilutions had OD_600_ ranging from 3.0, 1.0, 0.5, 0.25, 0.125 and 0.015625. Life span summary is reported in **Table S1.** Statistical analysis was performed using Two-way Annova.

#### Life span assay with 2-Deoxyglucose (2-DOG)

Plates for performing 2-DOG life span were prepared from the same batch of NGM agar as the control plates except that the 2-DOG (Sigma-Aldrich, D8375) was added in the media to a final concentration of 5 mM from a sterile 0.5 M stock solution made in water. Synchronized egg population (100-150) obtained by sodium hypochlorite treatment of gravid worms grown on *E. coli* OP50, was exposed to control plates (without 2-DOG). At YA stage, approximately 50 worms were transferred to the plates containing 2-DOG and plates without 2-DOG, overlaid with FUDR, in three technical replicates. Worms were scored every alternate day as mentioned above.

#### Dauer assay

Gravid adult worms were bleached and eggs were kept on OP50-seeded plates and upshifted to 22.5°C in an incubator. After 72h, worms were counted as adults or dauers. Percentage dauers for each strain was calculated and plotted. Two biological repeats, each with at least two technical repeats, were used for averaging.

#### RNA isolation, cDNA synthesis, and quantitative real time PCR

Eggs were synchronized by overnight starvation. These L1s were then allowed to grow till YA stage. YA worms were collected in M9 buffer after washing thrice to get rid of bacteria and then frozen in Trizol (Invitrogen, USA). The frozen worms were passed through two freeze-thaw cycles and lysed by vigorous vortexing. RNA was extracted by phenol:chloroform:isoamyl alcohol followed by ethanol precipitation. RNA was dissolved in DEPC-treated MQ water and denatured at 65°C for 10 min. The concentration of the RNA was then determined using NanoDrop 2000 (Thermo Scientific, USA). The integrity of the RNA was checked by electrophoresis on a denaturing formaldehyde gel.

First strand cDNA was synthesized using 2.5 μg RNA, employing Superscript III Reverse Transcriptase (Invitrogen, USA). Quantitative real time PCR (qRT-PCR) reaction was set up using DyNAmo Flash SYBR Green master mix (Thermo Scientific, USA) in a Realplex PCR system (Eppendorf, USA), according to manufacturer’s specifications. The gene expression was represented as relative fold change determined after normalizing the ΔCt values to *actin*, the housekeeping gene. Unpaired two-tailed *t*-test was applied as statistical analysis by using GraphPad Prism (GraphPad Software, La Jolla California).

#### Autophagosome quantification

Gravid adult worms expressing *lgg-1::gfp* were bleached and eggs were hatched on OP50 or RNAi bacteria. When they reach the L3 larval stage, worms were anesthetised with 20 mM sodium azide on 2% agarose slides and imaged at 630X using an AxioImager M2 microscope fitted with Axiocam MRc camera (Carl Zeiss, Germany). The autophagosomes were manually scored as GFP puncta in the hypodermal seam cells, seen only at L3stage. At least 15 worms were examined for approximately 3-10 seam cells in each one of them. The total number of puncta per seam cells for each worm was calculated and averaged out. Assay was repeated two times.

#### Western Blotting

Ten micrograms of protein for each sample was resolved on a 15% SDS-PAGE, transferred onto a PVDF membrane (Millipore, Billerica, MA) and blocked with 5% w/v skimmed milk protein prepared in 0.1% TBST (room temperature, 1 hr). Subsequently, the membrane was incubated overnight with anti-GFP antibody (Novus, USA - Cat. No. NB100-2220; 1:1000 dilution in 0.1% TBST) at 4 °C on a rocker-shaker. Following three washes of 5 minutes each in 0.1% TBST, anti-mouse secondary antibody (Abcam, UK - Cat. No. ab6728; 1:5000 dilution in 0.1% TBST) was added to the membrane and incubated for 1 hour at room temperature, with rocking. The membrane was then washed three times with 0.1% TBST, each for 5 minutes, at RT on a rocker-shaker. The blot was developed using an ECL reagent (Millipore, USA), according to manufacturer’s instructions.

#### Quantification of fat content by Oil Red O staining

Fat content of worms was determined according to previously published protocols (Chamoli et al. 2014; O’Rourke et al. 2009). Prepared in advance (Oil Red O working solution): Oil Red O stain was prepared as 5mg/ml stock in isopropanol and equilibrated on a rocker shaker for a week. The working stock of Oil Red O was prepared by diluting equilibrated stock to 60% with water. Stock was mixed thoroughly and filtered using a 0.22μm filter to remove any particles.

Eggs extracted from gravid adults by sodium hypochlorite treatment were grown till L4 or YA stage. The worms were washed in 1X PBS to remove any attached bacteria and resuspended in 120μl 1X PBS. To this an equal volume of 2X MRWB buffer (160mM KCl, 40mM NaCl, 14 mMNa2EGTA, PIPES pH 7.4, 1mM Spermidine, 0.4mM Spermine, 2% Paraformaldehyde, 0.2% β-mercaptoethanol) was added and incubated for 45 minutes on a rocker shaker. The worms were stored at −80 °C after slow freezing using liquid nitrogen. For staining, the frozen worms were thawed on ice, then pelleted and washed thrice with 1XPBS. An 120μl aliquot of the working solution of Oil Red O was added to the fixed worms and incubated for an hour on a shaker at room temperature. Following staining, worms were washed thrice with 1X PBS and mounted on 2% agarose slides for visualization using an AxioImager M2 microscope fitted with Axiocam MRc camera. ImageJ software was used to quantify fat stores in the intestine of worms.

#### Statistical analysis

Unless mentioned in the figure legends, at least three independent repeats were performed for all the experiments. All *p*-values were calculated (by two-tailed *t*-test or two-way Annova) and graphs were plotted using Graph pad Prism. For life span analysis, Mantel-Cox log rank test using OASIS software available at http://sbi.postech.ac.kr/oasis (Han et al. 2016) was used.

## Acknowledgements

We would like to thank the present and former members of the Molecular Aging lab for their scientific inputs. The research was supported in part by the National Bioscience Award for Career Development (BT/HRD/NBA/38/04/2016) and SERB-STAR award (STR/2019/000064) to AM, DBT’s Twinning Programme for the NE (BT/PR16823/NER/95/304/2015) to AM and AB, and core funding from the National Institute of Immunology to AM. Some strains were provided by the Caenorhabditis Genetics Center, which is funded by National Institute of Health Office of Research Infrastructure Programs (P40 OD010440).

---

**Table S1:** Summary of life span analysis, related to Figures 1, 2, 6 and Figures S4

**Table S2:** List of primers used in the study, related to Figure 3 and Figures 5.

## Ethics approval

(include appropriate approvals or waivers)

Not applicable

## Consent to participate

(include appropriate statements)

Not applicable

## Consent for publication

(include appropriate statements)

All authors consent to the publication of the article

## Availability of data and material

(data transparency)

All data is available on request

## Code availability

(software application or custom code)

Not applicable

## Authors’ contributions

(optional: please review the submission guidelines from the journal whether statements are mandatory)

AM conceived and supervised the project. AM and AB acquired funds and administered the project. AG performed all the experiments and analysed data. AG wrote the manuscript with AM, AB edited it.

---

**Figure S1.**
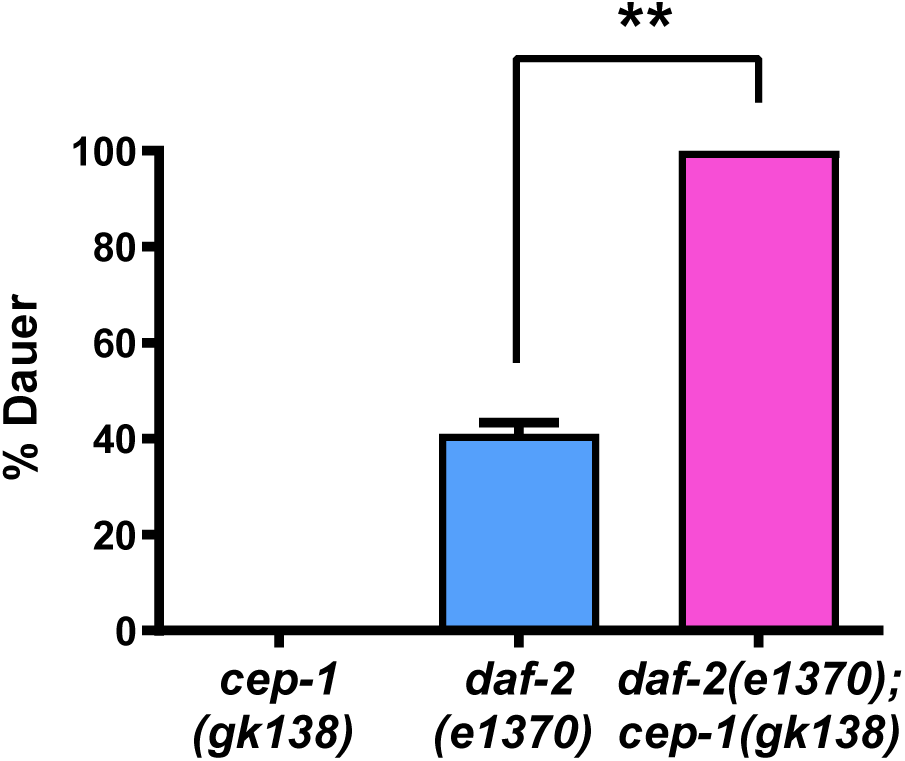
The percentage of dauer formed in *daf-2(e1370)* is enhanced in the genetic double *daf-2(e1370);cep-1(gk138)*. Dauer assay was performed at 22.5 °C. Average of two biological replicates is shown Error bars represent SEM. Unpaired two-tailed *t*-test. \*\**P*≤0.01.

**Figure S2.**
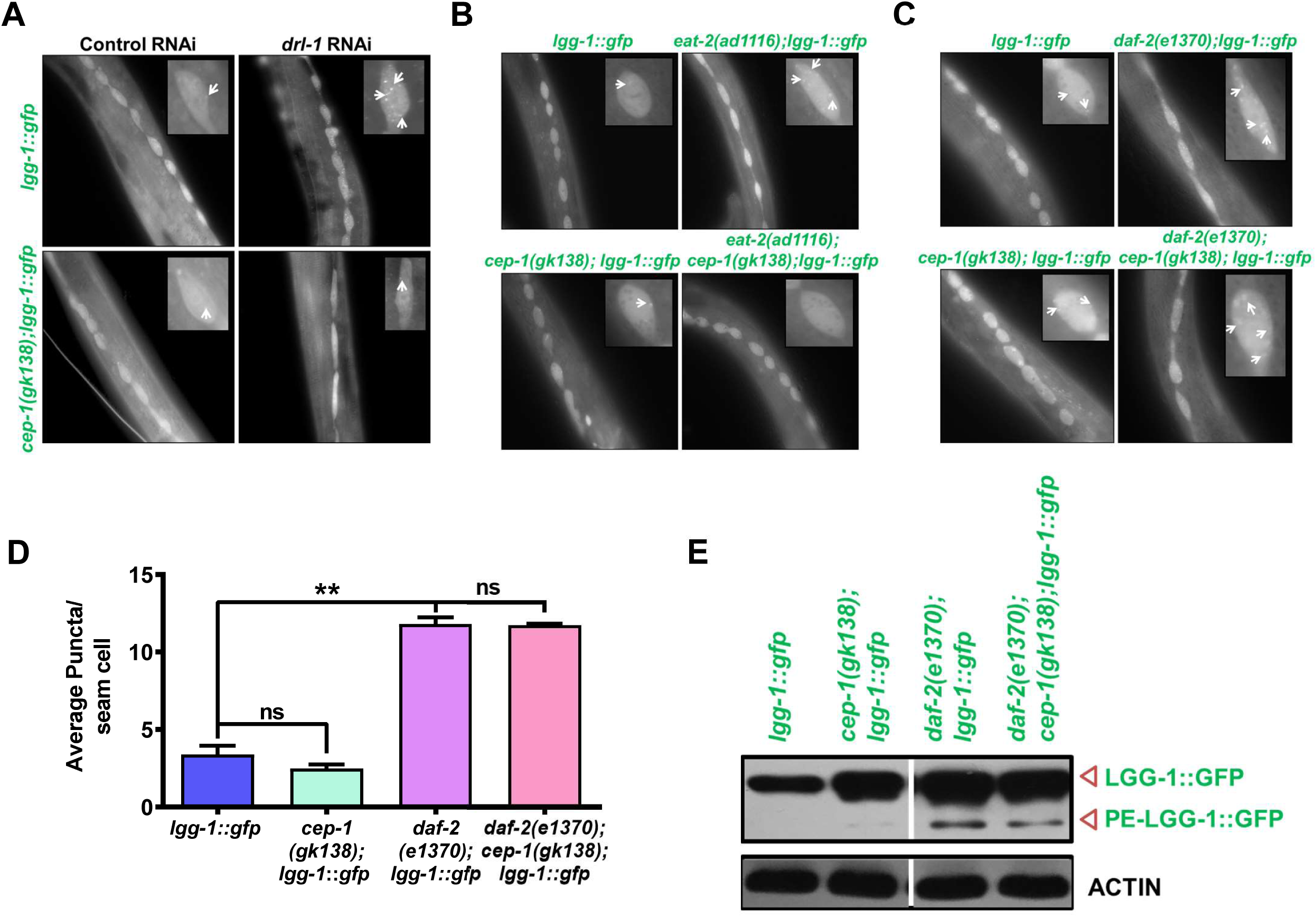
Representative images showing autophagosomes as GFP puncta in the hypodermal seam cells of L3-staged **(A)** *lgg::gfp* and *cep-1(gk138)*;*lgg::gfp* worms on control or *drl-1* RNAi, **(B)** *eat-2(ad116);lgg-1::gfp* and *eat-2(ad116);cep-1(gk138)*;*lgg-1:gfp*, **(C)** *daf-2(e1370);lgg-1::gfp* and *daf-2(e1370);cep-1(gk138)*;*lgg-1::gfp*. Images were captured at 630X magnification. Inset images represent zoomed-in areas showing one seam cell. Arrows point to autophagosome puncta. **(D)** The increased autophagosome formation in *daf-2(e1370);lgg-1:gfp* is unaffected in *daf-2(e1370);cep-1(gk138)*;*lgg-1::gfp.* Quantification of GFP puncta averaged over 2 biological repeats is shown. Error bars represent SEM. No. of animals analysed n≥15. Unpaired two-tailed t-test. \*\**P*≤0.01, ns= non-significant. **(E)** Western blot using anti-GFP antibody to detect the unmodified LGG-1 or PE-LGG-1 in *daf-2(e1370);lgg-1::gfp* and *daf-2(e1370);cep-1(gk138)*;*lgg-1::gfp*. β-ACTIN was used as a loading control. One representative blot out of four biological replicates is shown.

**Figure S3.**
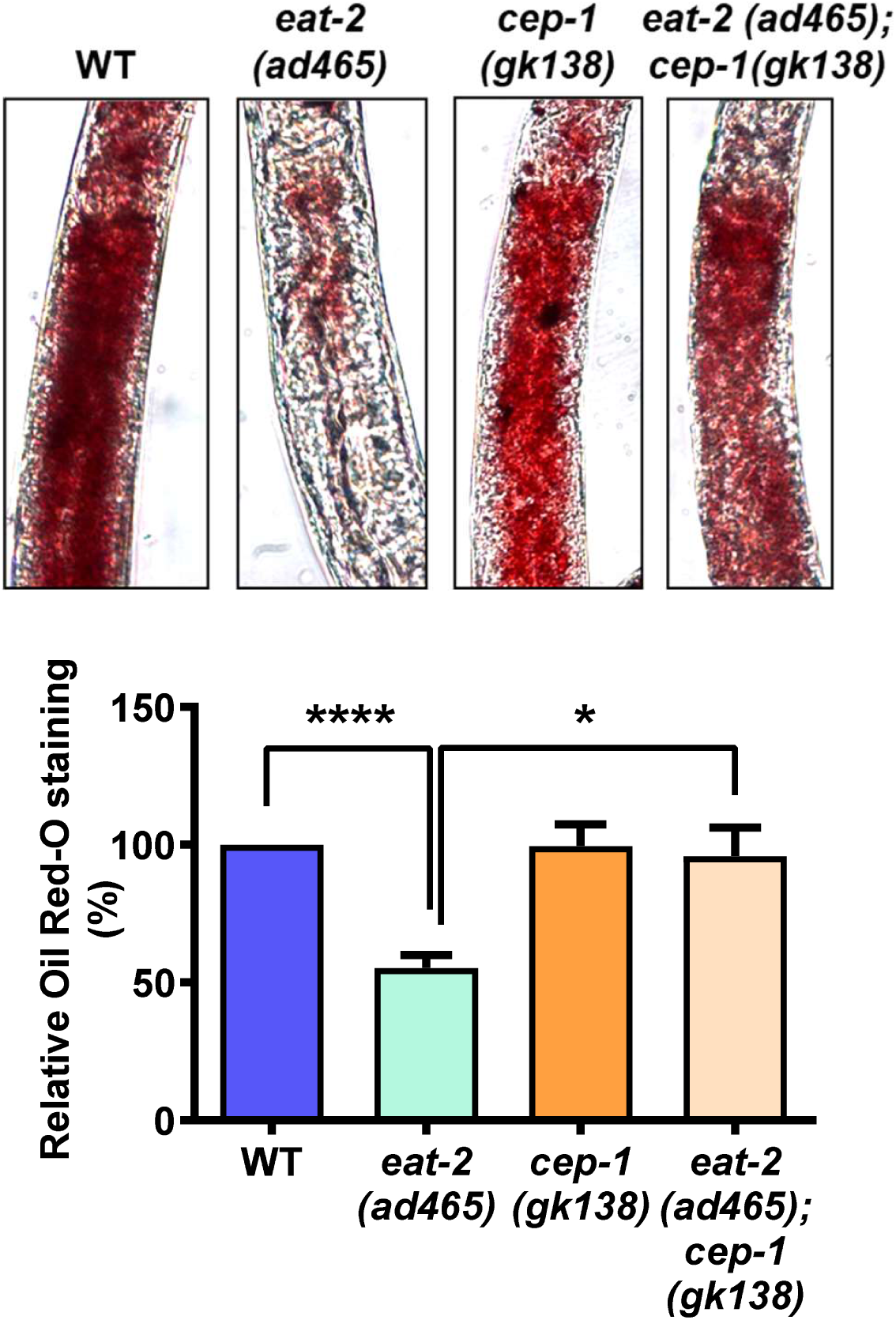
Oil Red O staining of *eat-2(ad465)* or *eat-2(ad465);cep-1(gk138)*. Lowering of fat storage in *eat-2(ad1116)* is attenuated in *eat-2(ad465);cep-1(gk138)*. Images were captured at 100X magnification. Quantification averaged over at least three biological replicates is shown below in each case. Error bars represent SEM. Unpaired two-tailed *t*-test. \**P*≤0.05, \*\*\*\**P*≤0.0001.

**Figure S4.**
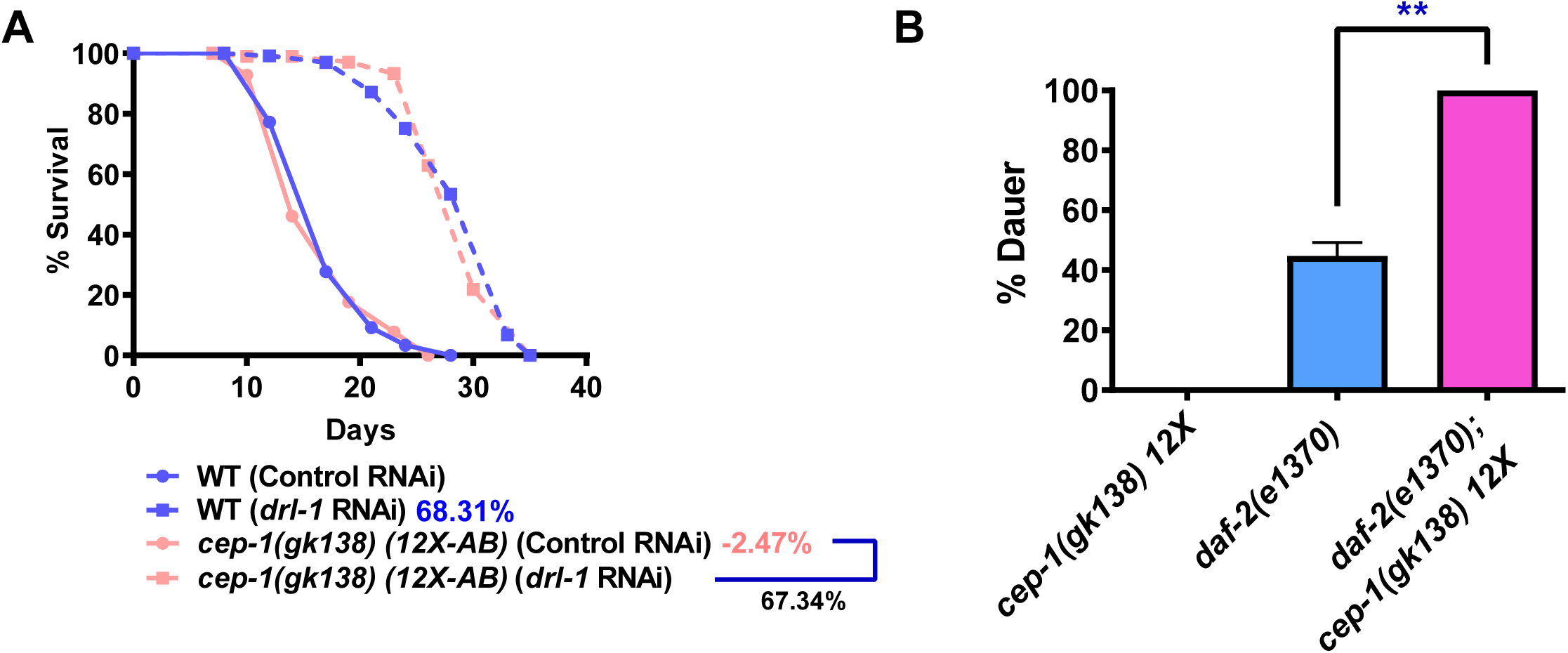
**(A)** Life span analysis of *cep-1(gk138)* (12X-AB) generated earlier (Baruah et al. 2014) as well as WT, grown on control or *drl-1* RNAi. Life span was performed at 20^°^C and details are provided in Table S1. **(B)** The percentage of dauer formed in *daf-2(e1370)* is enhanced in the genetic double *daf-2(e1370);cep-1(gk138) 12X*. Dauer assay was performed at 22.5 °C. Average of three biological replicates is shown. Error bars represent SEM. Unpaired two-tailed *t*-test. \*\**P*≤0.01.

