## Supplemental Table 1 for "The genetic paradigms of dietary restriction fail to extend life span in *cep-1(gk138)* mutant of *C. elegans* p53 due to possible background mutations"

| **Table S1: Summary of life span analysis, related to Figures 1, 2, 6 and Figures S4** | | | | | | | | |
|---|---|---|---|---|---|---|---|---|
| **Expt. No.** | **Genetic Background** | **RNAi** | **Mean ± SEM** | **% change w.r.t. WT on control RNAi** | **% change w.r.t. *cep-1(gk138)* control RNAi** | **n (Number of animals)** | ***P*-value w.r.t. WT control RNAi** | ***P*-value w.r.t. *cep-1(gk138)* control RNAi** |
| 1 | WT **(Figure 1A)** | control | 20.19 ± 0.37 | | | 118 | | |
| | | *drl-1* | 31.09 ± 0.55 | (+) 53.99 | | 87 | ≤0.0001 | |
| | *cep-1(gk138)* | control | 22.63 ± 0.23 | (+) 12.08 | | 124 | ≤0.0001 | |
| | | *drl-1* | 22.68 ± 0.56 | | (+) 0.22 | 78 | | 0.1981 |
| 2 | WT | control | 18.78 ± 0.36 | | | 83 | | |
| | | *drl-1* | 25.58 ± 0.44 | (+) 36.20 | | 72 | ≤0.0001 | |
| | *cep-1(gk138)* | control | 23.31 ± 0.15 | (+) 24.12 | | 180 | ≤0.0001 | |
| | | *drl-1* | 22.70 ± 0.57 | | (-) 2.61 | 79 | | 0.7831 |
| **Expt. No.** | **Genetic Background** | **Bacteria** | **Mean ± SEM** | **% change w.r.t. WT** | **% change w.r.t. *cep-1(gk138)*** | **n (Number of animals)** | ***P*-value w.r.t. WT** | ***P*-value w.r.t. *cep-1(gk138)*** |
| 1 | WT **(Figure 1B)** | OP50 | 18.64 ± 0.23 | | | 194 | | |
| | *cep-1(gk138)* | OP50 | 20.49 ± 0.16 | (+) 09.92 | | 226 | ≤0.0001 | |
| | *eat-2(ad1116)* | OP50 | 26.43 ± 0.58 | (+) 41.79 | | 79 | ≤0.0001 | |
| | *eat-2(ad1116); cep-1(gk138)* | OP50 | 20.28 ± 0.37 | | (-) 1.02 | 158 | | 0.9533 |
| 2 | WT | OP50 | 18.42 ± 0.29 | | | 145 | | |
| | *cep-1(gk138)* | OP50 | 20.65 ± 0.28 | (+) 12.11 | | 167 | ≤0.0001 | |
| | *eat-2(ad1116)* | OP50 | 27.36 ± 0.40 | (+) 48.53 | | 83 | ≤0.0001 | |
| | *eat-2(ad1116); cep-1(gk138)* | OP50 | 21.45 ± 0.26 | | (+) 3.87 | 144 | | 0.6353 |
| 1 | WT **(Figure 1C)** | OP50 | 18.01 ± 0.19 | | | 175 | | |
| | *cep-1(gk138)* | OP50 | 20.63 ± 0.22 | (+) 14.54 | | 167 | ≤0.0001 | |
| | *eat-2(ad465)* | OP50 | 23.35 ± 0.32 | (+) 29.65 | | 142 | ≤0.0001 | |
| | *eat-2(ad465); cep-1(gk138)* | OP50 | 19.77 ± 0.29 | | (-) 4.16 | 208 | | 0.3199 |
| 2 | WT | OP50 | 18.42 ± 0.29 | | | 145 | | |
| | *cep-1(gk138)* | OP50 | 20.65 ± 0.28 | (+) 12.11 | | 167 | ≤0.0001 | |
| | *eat-2(ad465)* | OP50 | 24.05 ± 0.24 | (+) 30.56 | | 133 | ≤0.0001 | |
| | *eat-2(ad465); cep-1(gk138)* | OP50 | 21.69 ± 0.27 | | (+) 5.03 | 169 | | 0.2706 |

| Expt. No. | Genetic Background | Bacteria | Mean ± SEM | % change w.r.t. WT | % change w.r.t. *cep-1(gk138)* | n (Number of animals) | *P*-value w.r.t. WT | *P*-value w.r.t. *cep-1(gk138)* |
|---|---|---|---|---|---|---|---|---|
| 1 | WT (**Figure 1D)** | OP50 | 21.33 ± 0.27 | | | 64 | | |
| | *cep-1(gk138)* | OP50 | 26.25 ± 0.40 | (+) 23.06 | | 118 | ≤0.0001 | |
| | *daf-2(e1370)* | OP50 | 45.25 ± 0.58 | (+) 112.14 | | 161 | ≤0.0001 | |
| | *daf-2(e1370); cep-1(gk138)* | OP50 | 42.81 ± 0.57 | | (+) 63.08 | 134 | | ≤0.0001 |
| 2 | WT | OP50 | 19.45 ± 0.24 | | | 80 | | |
| | *cep-1(gk138)* | OP50 | 22.15 ± 0.41 | (+)13.88 | | 62 | ≤0.0001 | |
| | *daf-2(e1370)* | OP50 | 39.38 ± 1.12 | (+)102.4 | | 74 | ≤0.0001 | |
| | *daf-2(e1370); cep-1(gk138)* | OP50 | 37.41 ± 1.43 | | (+) 68.89 | 70 | | ≤0.0001 |
| **Expt. No.** | **Genetic Background** | **Dilution** | **Mean ± SEM** | **% change** | | **n (Number of animals)** | ***P*-value** | |
| 1 | WT **(Figure 2A)** | Dil 1 (OD 3.0) | 30.46± 0.51 | | | 52 | | |
| | | Dil 2 (OD 1.0) | 41.15 ± 0.99 | (+) 35.09 | | 46 | ≤0.0001 | |
| | | Dil 3 (OD 0.5) | 43.08 ± 0.83 | (+) 30.66 | | 48 | ≤0.0001 | |
| | | Dil 4 (OD 0.25) | 42.27 ± 0.93 | (+) 27.41 | | 45 | ≤0.0001 | |
| | | Dil 5 (OD 0.125) | 37.06 ± 1.07 | (+) 15.61 | | 47 | ≤0.0001 | |
| | | Dil 6 (OD 0.015) | 37.17 ± 0.81 | (+) 18.10 | | 41 | ≤0.0001 | |
| | *cep-1(gk138)* | Dil 1 (OD 3.0) | 30.7 ± 0.52 | | | 46 | | |
| | | Dil 2 (OD 1.0) | 40.78 ± 0.64 | (+) 32.83 | | 41 | ≤0.0001 | |
| | | Dil 3 (OD 0.5) | 44.67 ± 0.67 | (+) 34.25 | | 42 | ≤0.0001 | |
| | | Dil 4 (OD 0.25) | 45.00 ± 0.62 | (+) 32.01 | | 46 | ≤0.0001 | |
| | | Dil 5 (OD 0.125) | 41.14 ± 0.71 | (+) 23.20 | | 43 | ≤0.0001 | |
| | | Dil 6 (OD 0.015) | 36.35 ± 1.14 | (+) 13.73 | | 46 | ≤0.0001 | |

| | | | | | | | |
|---|---|---|---|---|---|---|---|
| 2 | WT | Dil 1 (OD 3.0) | 28.23 ± 0.65 | | | 44 | |
| | | Dil 2 (OD 1.0) | 44.37 ± 0.73 | (+) 57.17 | | 46 | ≤0.0001 |
| | | Dil 3 (OD 0.5) | 48.36 ± 0.83 | (+) 45.36 | | 45 | ≤0.0001 |
| | | Dil 4 (OD 0.25) | 36.02 ± 0.93 | (+) 16.10 | | 41 | ≤0.0001 |
| | | Dil 5 (OD 0.125) | 35.13 ± 1.07 | (+) 19.15 | | 40 | ≤0.0001 |
| | | Dil 6 (OD 0.015) | 32.13 ± 0.81 | (+) 11.10 | | 40 | ≤0.0001 |
| | *cep-1(gk138)* | Dil 1 (OD 3.0) | 29.70 ± 0.79 | | | 46 | |
| | | Dil 2 (OD 1.0) | 37.40 ± 1.28 | (+) 25.92 | | 48 | ≤0.0001 |
| | | Dil 3 (OD 0.5) | 44.02 ± 1.54 | (+) 38.28 | | 49 | ≤0.0001 |
| | | Dil 4 (OD 0.25) | 37.54 ± 0.85 | (+) 17.81 | | 46 | ≤0.0001 |
| | | Dil 5 (OD 0.125) | 33.34 ± 1.40 | (+) 09.69 | | 47 | ≤0.0001 |
| | | Dil 6 (OD 0.015) | 32.47 ± 1.67 | (+) 08.30 | | 45 | ≤0.0001 |

| **Expt. No.** | **Genetic Background** | **+/- DOG** | **Mean ± SEM** | **% change w.r.t. WT** | **% change w.r.t. *cep-1(gk138)*** | **n (Number of animals)** | ***P*-value w.r.t. WT** | ***P*-value w.r.t. *cep-1(gk138)*** |
|---|---|---|---|---|---|---|---|---|
| 1 | WT **(Figure 2B)** | OP50 | 18.64 ± 0.23 | | | 194 | | |
| | | 2-DOG + OP50 | 23.32 ± 0.36 | (+) 25.10 | | 153 | ≤0.0001 | |
| | *cep-1(gk138)* | OP50 | 20.49 ± 0.16 | (+) 09.92 | | 226 | ≤0.0001 | |
| | | 2-DOG + OP50 | 26.37 ± 0.31 | | (+) 28.69 | 145 | | ≤0.0001 |
| 2 | WT | OP50 | 18.42 ± 0.29 | | | 145 | | |
| | | 2-DOG + OP50 | 20.35 ± 0.51 | (+) 10.47 | | 71 | 0.0010 | |
| | *cep-1(gk138)* | OP50 | 20.65 ± 0.28 | (+) 12.10 | | 167 | ≤0.0001 | |
| | | 2-DOG + OP50 | 24.49 ± 0.28 | | (+) 18.59 | 148 | | ≤0.0001 |
| 1 | WT **(Figure 6A)** | control | 18.78 ± 0.36 | | | 83 | | |
| | | *drl-1* | 25.58 ± 0.44 | (+) 36.20 | | 72 | ≤0.0001 | |

| | | | | | | | | |
|---|---|---|---|---|---|---|---|---|
| | *cep-1(gk138) 11X* | control | 23.01 ± 0.37 | (+) 22.52 | | 88 | ≤0.0001 | |
| | | *drl-1* | 21.92 ± 0.40 | | (-) 4.73 | 92 | | 0.0746 |
| 2 | WT | control | 19.79 ± 0.27 | | | 98 | | |
| | | *drl-1* | 30.14 ± 0.65 | (+) 52.29 | | 63 | ≤0.0001 | |
| | *cep-1(gk138) 11X* | control | 23.92 ± 0.40 | (+) 20.86 | | 90 | ≤0.0001 | |
| | | *drl-1* | 23.92 ± 0.71 | | 0 | 74 | | 0.3452 |
| 1 | WT **(Figure 6B)** | control | 17.39 ± 0.36 | | | 119 | | |
| | | *drl-1* | 29.27 ± 0.43 | (+) 68.31 | | 133 | ≤0.0001 | |
| | *cep-1(gk138) 12X* | control | 16.23 ± 0.43 | (-) 06.67 | | 129 | 0.1374 | |
| | | *drl-1* | 27.16 ± 0.45 | | (+) 67.34 | 117 | | ≤0.0001 |
| 2 | WT | control | 21.13 ± 0.40 | | | 141 | | |
| | | *drl-1* | 32.27 ± 0.31 | (+) 52.72 | | 147 | ≤0.0001 | |
| | *cep-1(gk138) 12X* | control | 22.58 ± 0.67 | (+) 06.86 | | 92 | 0.0173 | |
| | | *drl-1* | 30.32 ± 0.33 | | (+) 34.27 | 141 | | ≤0.0001 |
| 1 | WT **(Figure 6C)** | control | 19.30 ± 0.37 | | | 98 | | |
| | | *drl-1* | 28.27 ± 0.24 | (+) 46.47 | | 194 | ≤0.0001 | |
| | *TJ1 [cep-1(gk138)]* | control | 22.57 ± 0.36 | (+) 16.94 | | 47 | ≤0.0001 | |
| | | *drl-1* | 21.96 ± 0.42 | | (-) 2.70 | 90 | | 0.6831 |
| 2 | WT | control | 19.92 ± 0.29 | | | 190 | | |
| | | *drl-1* | 25.43 ± 0.23 | (+) 27.66 | | 188 | ≤0.0001 | |
| | *TJ1 [cep-1(gk138)]* | control | 23.70 ± 0.39 | (+) 18.97 | | 98 | ≤0.0001 | |
| | | *drl-1* | 21.66 ± 0.40 | | (-) 8.60 | 108 | | 0.0021 |

| | | | | | | | | |
|---|---|---|---|---|---|---|---|---|
| 1 | WT **(Figure 6D)** | control | 19.30 ± 0.37 | | | 98 | | |
| | | *drl-1* | 28.27 ± 0.24 | (+) 46.47 | | 194 | ≤0.0001 | |
| | *cep-1(gk138) VC172* | control | 21.73 ± 0.41 | (+) 12.59 | | 70 | ≤0.0001 | |
| | | *drl-1* | 30.48 ± 0.53 | | (+) 40.26 | 63 | | ≤0.0001 |
| 2 | WT | control | 18.78 ± 0.36 | | | 83 | | |
| | | *drl-1* | 25.58 ± 0.44 | (+) 36.20 | | 72 | ≤0.0001 | |
| | *cep-1(gk138) VC172* | control | 23.43 ± 0.64 | (+) 24.76 | | 84 | ≤0.0001 | |
| | | *drl-1* | 29.79 ± 0.48 | | (+) 27.14 | 63 | | ≤0.0001 |

| | | | | | | | | |
|---|---|---|---|---|---|---|---|---|
| 1 | WT **(Figure 6F)** | control | 19.58 ± 0.28 | | | 84 | | |
| | | *drl-1* | 30.40 ± 0.47 | (+) 55.26 | | 85 | ≤0.0001 | |
| | *cep-1(ep347)* | control | 21.97 ± 0.48 | (+) 12.20 | | 59 | ≤0.0001 | |
| | | *drl-1* | 30.25 ± 0.39 | | (+) 37.68 | 91 | | ≤0.0001 |
| 2 | WT | control | 18.32 ± 0.36 | | | 91 | | |
| | | *drl-1* | 25.46 ± 0.28 | (+) 38.97 | | 127 | ≤0.0001 | |
| | *cep-1(ep347)* | control | 20.93 ± 0.50 | (+) 14.24 | | 57 | ≤0.0001 | |
| | | *drl-1* | 28.25 ± 0.29 | | (+) 34.97 | 178 | | ≤0.0001 |
| 1 | WT **(Figure 6E)** | control | 19.58 ± 0.28 | | | 84 | | |
| | | *drl-1* | 30.40 ± 0.47 | (+) 55.26 | | 85 | ≤0.0001 | |
| | *cep-1(lg12501)* | control | 23.11 ± 0.23 | (+) 18.02 | | 70 | ≤0.0001 | |
| | | *drl-1* | 31.07 ± 0.73 | | (+) 34.44 | 54 | | ≤0.0001 |
| 2 | WT | control | 18.32 ± 0.36 | | | 91 | | |
| | | *drl-1* | 25.46 ± 0.28 | (+) 38.97 | | 127 | ≤0.0001 | |
| | *cep-1(lg12501)* | control | 22.42 ± 0.44 | (+) 22.37 | | 52 | ≤0.0001 | |
| | | *drl-1* | 26.14 ± 0.36 | | (+) 16.59 | 90 | | ≤0.0001 |
| 1 | WT **(Figure S4A)** | control | 17.39 ± 0.36 | | | 119 | | |
| | | *drl-1* | 29.27 ± 0.43 | (+) 68.31 | | 133 | ≤0.0001 | |
| | *cep-1(gk138) 12X-AB* | control | 16.96 ± 0.36 | (-) 02.47 | | 141 | 0.885 | |
| | | *drl-1* | 21.21 ± 0.41 | | (+) 72.22 | 105 | | ≤0.0001 |
| 2 | WT | control | 21.13 ± 0.40 | | | 141 | | |
| | | *drl-1* | 32.27 ± 0.31 | (+) 52.72 | | 147 | ≤0.0001 | |
| | *cep-1(gk138) 12X-AB* | control | 21.07 ± 0.56 | (-) 00.28 | | 97 | 0.8333 | |
| | | *drl-1* | 32.50 ± 0.40 | | (+) 54.24 | 116 | | ≤0.0001 |

*w.r.t. = with respect to.
