## Supplemental Table 2 for "The genetic paradigms of dietary restriction fail to extend life span in *cep-1(gk138)* mutant of *C. elegans* p53 due to possible background mutations"

**Table 2: List of qRT-PCR primers used**

| Gene Name (Target) | Primer name | Sequence |
|---|---|---|
| *lgg-1* | Forward | GGT CCC ATC CGA TCT TAC TGT |
| *lgg-1* | Reverse | CGT GAT GGT CCT GGT AGA GT |
| *bec-1* | Forward | GAA AGA GCT CAA GGA TCG AAA CA |
| *bec-1* | Reverse | GAG CAT TAG AGC CAT TGC ACG |
| *unc-51* | Forward | CGC TAT GTT GAT CGC ACA GAC |
| *unc-51* | Reverse | GTG CAT TTG AGT AGG CCC AC |
| *vps-34* | Forward | CCT TAT TTG GTT CTC TCC ACA G |
| *vps-34* | Reverse | CCA TCT CCT GGA CGA AGT TC |
| *ugt-16* | Forward | CTTGCTGACGATCGACTAACC |
| *ugt-16* | Reverse | CGGTCTGTATGGCTTCTCTAAG |
| *gst-6* | Forward | CAAAAATAACACTCCATTC |
| *gst-6* | Reverse | GCCGCCTCGGTGTCATTTTGTC |
| *gst-28* | Forward | GCTTAAAGACGGCGCCCC |
| *gst-28* | Reverse | CCAGCATATCCGAATTTGTTGGC |
| *cyp-32B1* | Forward | GGTGTGTTGAAGTTATGGTTGGGACC |
| *cyp-32B1* | Reverse | TGTCGCCGGTGCTGATTAAAAGAC |
| *cyp-33C8* | Forward | CGCTGGATGATGTGCTCAACTACTGG |
| *cyp-33C8* | Reverse | GCTTCTTCTGCTCTTTCAGGTAGG |
| *cyp-34A4* | Forward | GATTTGAACAGGGTGACCCAGAAT |
| *cyp-34A4* | Reverse | TCGATGACATGCTCACCACT |
| *cyp-37B1* | Forward | GCTTGGAACGGGACTATTGAC |
| *cyp-37B1* | Reverse | TTGTTCGAGGAAAACCTTGGCCTG |
| *actin* | Forward | CTCTTGCCCCATCAACCATG |
| *actin* | Reverse | CTTGCTTGGAGATCCACATC |
